# Feature-based attention enables robust, long-lasting location transfer in human perceptual learning

**DOI:** 10.1101/2021.04.28.441788

**Authors:** Shao-Chin Hung, Marisa Carrasco

## Abstract

Visual perceptual learning (VPL) is typically specific to the trained location and feature. However, the degree of specificity depends upon particular training protocols. Manipulating covert spatial attention during training facilitates learning transfer to other locations. Here we investigated whether feature-based attention (FBA), which enhances the representation of particular features throughout the visual field, facilitates VPL transfer, and how long such an effect would last. To do so, we implemented a novel task in which observers discriminated a stimulus orientation relative to two reference angles presented simultaneously before each block. We found that training with FBA enabled remarkable location transfer, reminiscent of its global effect across the visual field, but preserved orientation specificity in VPL. Critically, both the perceptual improvement and location transfer persisted after one year. Our results reveal robust, long-lasting benefits induced by FBA in VPL, and have translational implications for improving generalization of training protocols in visual rehabilitation.

## Introduction

The available sensory information at any given moment is far too much for the visual system to process at once. To function effectively, the visual system must establish stability and selectively process the most important information. In the short term, visual attention allows us to select relevant visual information; in the long term, perceptual learning allows us to adapt to new surroundings and to refine how sensory systems efficiently process stimuli that we regularly experience.

Perceptual learning refers to improvements in sensory discrimination due to repetitive practice, which has been observed in multiple sensory modalities, including visual [1], auditory [2], tactile [3] and olfactory [4]. Perceptual learning is considered a manifestation of neural plasticity in the adult brain, enabling adaptive responses to environmental changes. Visual perceptual learning (VPL) has been demonstrated in various basic visual dimensions, such as orientation [5–7], contrast [8–10], texture [1,11,12], hyperacuity [13–16] and motion direction [17–21]. VPL often requires thousands of trials of practice over days or weeks, and performance improvement can last for months or even years [9,22–26].

A hallmark of VPL is that learning is typically highly specific to the trained location and feature (e.g. orientation, motion direction) [1,6,14–16,18,19,27,28]. This specificity is typically interpreted as evidence that VPL occurs in early cortical regions [7,29–32], where receptive fields of neurons are selective for these attributes.

The degree of learning specificity, however, depends upon specifics of the experimental procedure; e.g.: length of training [33], difficulty of training stimuli [34,35], inclusion of a pretest [36], whether multiple stimuli are trained [35,37,38], adaptation [11], and deployment of covert spatial attention [5,27,39,40]. For example, a double-training protocol enables transfer of learning to a different location by employing a secondary task [37]. Investigating factors and protocols that influence learning specificity and transfer provides a theoretical framework to infer cortical plasticity underlying VPL. Compared to specificity as a typical training outcome, transfer underscores the potential for translating VPL into a systematic training regime to improve visual skills and rehabilitation. An efficient training regime should promote learning to untrained conditions to maximize training benefits. Thus, understanding when and why training leads to transfer has become a central focus in VPL.

Recent research has highlighted the importance of top-down attentional modulation in VPL. The role of attention – the process by which rich sensory information is selected and prioritized – has been often discussed [41–44] but rarely manipulated and isolated in VPL. Thus, it is still largely unknown how attention and VPL interact and whether and how their underlying mechanisms are related.

To date, only a few studies have explicitly isolated attentional effects on VPL specificity. Research in our laboratory has revealed that both exogenous (involuntary) and endogenous (voluntary) spatial attention [45] during training facilitate transfer of learning to untrained locations [5,27,40]. Of note, manipulating attention during a single task requires less time and effort than other protocols that employ a secondary task to induce transfer [37,38,46], and is thus a more efficient training regime. Therefore, characterizing the effects of distinct types of attention on VPL will inform the development of efficient training protocols and shed light on how VPL relies on plasticity across different brain areas.

Feature-based attention (FBA), the selective processing of a relevant feature over unattended features, is notable in terms of its “global” effect. In contrast to spatial attention, which enhances processing within a spatial focus, behavioral and neuroimaging FBA studies have demonstrated its location-independent property: FBA is deployed simultaneously throughout the visual field, including locations that are irrelevant to the observer’s current task [47–58]. There is ample evidence characterizing FBA effects on visual perception, but it is unknown whether FBA generalizes VPL. To inform the development of efficient training protocols, and to gain mechanistic insight of VPL, we investigate whether and to which extent FBA influences the degree of specificity in human VPL.

To address this question, first we implemented a novel orientation discrimination task in which observers were presented with two reference angles simultaneously before each block, then asked to discriminate whether the orientation of a Gabor stimulus was clockwise or counter-clockwise with respect to either reference during each trial. In Experiment 1, we confirmed that FBA improves accuracy in this task.

Then we investigated the effects of FBA on location and feature specificity in VPL. In Experiment 2, two groups of observers participated in a six-day study; the Attention group trained with a feature attention cue, and the Neutral group trained with a neutral cue. To isolate the effects of training with FBA on VPL, observers were presented with a neutral cue during both the pre-test (before training) and post-test (after training) sessions. Because of the global effect of FBA on the attended features, we hypothesized that observers deploying feature attention during training would overcome retinotopic specificity, but not orientation specificity, whereas the Neutral group would exhibit both location and orientation specificity.

VPL improvement can last for months or even years [9,22–26]. It is unknown, however, how long the learning transfer to untrained conditions may last. To assess the duration of our observed VPL effects, we re-tested the observers 3-4 months, and ~1 year after training. We hypothesized that for both groups VPL at the trained location and orientation would be long lasting, and investigated whether any *transfer* effect would be long lasting.

Our results show perceptual benefits of FBA on an orientation discrimination task and reveal remarkable spatial-transfer in the Attention group, whereas the Neutral group exhibited both location and orientation specificity. Critically, the perceptual improvement attained by both groups and the location transfer attained by the Attention group were preserved for over a year. The robust and long-lasting training benefits enabled by FBA imply that it is a useful tool to potentiate the benefits of VPL by enabling generalization via location transfer. These novel results suggest an interaction between top-down FBA modulation and processing in visual cortices, thus providing converging evidence that VPL arises from plasticity across multiple cortical areas [59].

## Results

### Perceptual Benefit of FBA

In Experiment 1, we first validated the effects of FBA on an orientation discrimination task (**Fig 1A)**. This single-session experiment consisted of 800 trials, half preceded by a neutral cue and the other half by an attentional cue. For each neutral or attentional condition (blocked), there were 4 different conditions (i.e., stimuli on the left or right, and reference orientations of 30°/120° or 60°/150°, which were shown at the beginning of each block, but not during stimulus presentation). To obtain a psychometric function, we had five offsets either clockwise or counter-clockwise (2°, 4°, 6°, 8°, and 10°) from the reference angles.

**Fig 1.**
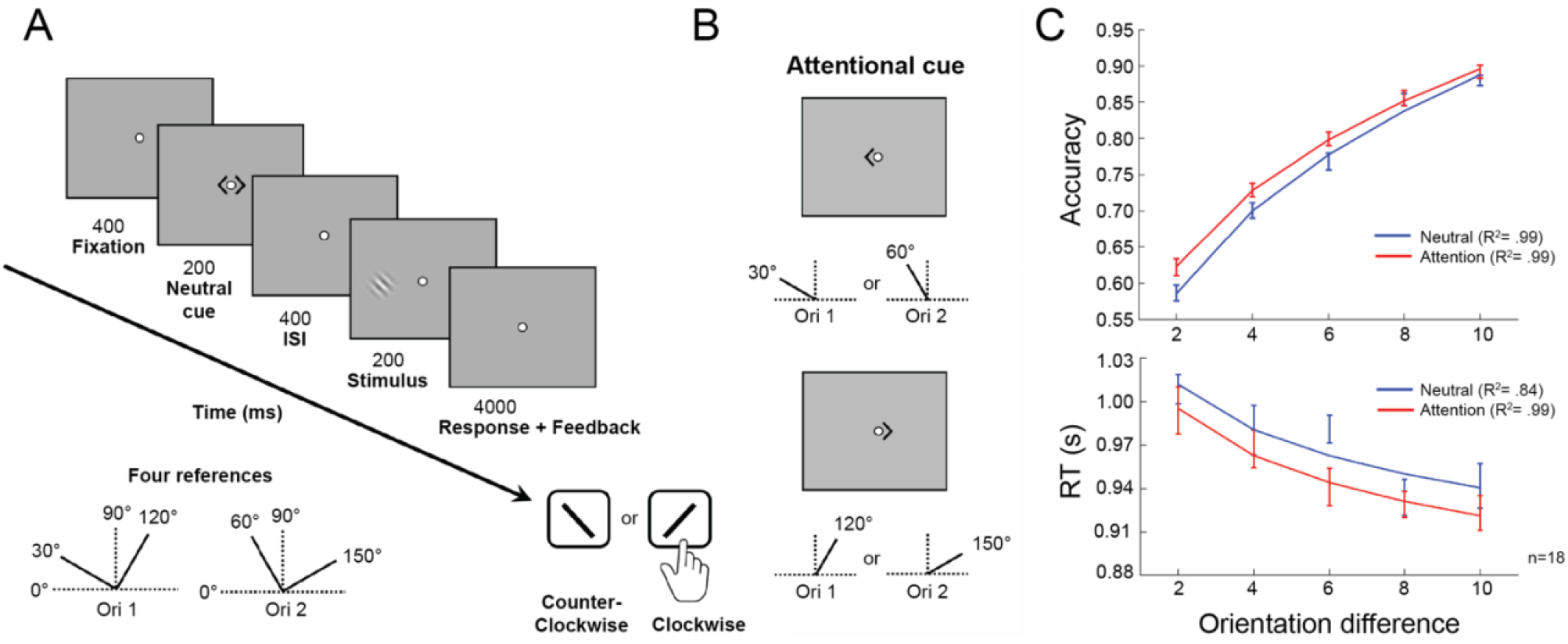
Perceptual benefit of feature-based attention in the orientation discrimination task. (**A**) Illustration of the orientation discrimination task. Before each block, one of the reference combinations (bottom left), each consisting of two reference orientations (30°/120°, or 60°/150°), was shown to observers, but never appeared on the screen during the stimulus presentation. In the task, each trial began with a fixation period of 400ms followed by a 200ms cue (neutral or attention). After 400ms ISI, a Gabor stimulus was presented for a single interval of 200ms, and the observer’s task was to judge whether the orientation of the stimulus was counter-clockwise or clockwise relative to the closest reference orientation shown before the block with a key-press within 4s. (**B**) Attentional cue. In the attention condition, the cue was either a leftward arrowhead indicating a reference angle of 30° or 60°, or a rightward arrowhead indicating 120° or 150°, depending on which reference combination was used in each block. (**C**) Deploying FBA (red line) significantly improved accuracy (upper panel) in the orientation discrimination task, without any speed-accuracy trade-off (lower panel) compared to the neutral condition (blue line). Error bars represent ±1 within-subject SEM.

In the attentional condition, participants were instructed to deploy their FBA to a particular reference orientation indicated by a cue before stimulus presentation (**Fig 1B**). A two-way repeated measures ANOVA was conducted to assess the effects of attention and offset sizes on accuracy. There were significant main effects of attention (*F*(1,17)=4.591, *p*=0.047) and offset size (*F*(4,68)=234.73, *p*<0.001), but no interaction between them (*F*(4,68)=1.253, *p*=0.297). That is, deploying FBA significantly increased discrimination accuracy across different offset sizes compared to the neutral condition (**Fig 1C, upper panel**, t_4_=3.428, *p*=0.027, Cohen’s d=0.191, two-tailed, paired t-test). We analyzed reaction time (RT) as a secondary measure to rule out a possible speed-accuracy trade-off in processing. A two-way repeated measures ANOVA revealed a significant main effect of offset sizes (*F*(4,68)=21.426, *p*<0.001), indicating that observers responded faster at larger offset sizes, when accuracy was higher (**Fig 1C, lower panel)**. Although RTs were overall faster in the attention condition than in the neutral condition (t_4_=3.106, *p*=0.036, Cohen’s d=0.605, two-tailed, paired t-test), neither the main effect of attention (*F*(1,17)<1) nor the interaction between attention and offset sizes (*F*(4,68)<1) was significant. In sum, this experiment confirmed that FBA enabled observers to perform more accurately, without any speed-accuracy trade-off, in this orientation discrimination task.

### Spatial-Transfer, but Not Feature-Transfer Induced by FBA in VPL

In Experiment 2, two groups of observers participated in a six-day study (**Figs 2A,B**). The Attention group trained with a feature attention cue, indicating which of the two reference angles was relevant for the discrimination on a trial-by-trial basis; the Neutral group trained with an uninformative neutral cue indicating both reference angles. Observers in both groups were presented with neutral cues during both the pre-test (before training) and post-test (after training) sessions. Given the global effect of FBA on the attended features, we hypothesized that observers in the Attention group would overcome retinotopic specificity but exhibit orientation specificity.

**Fig 2.**
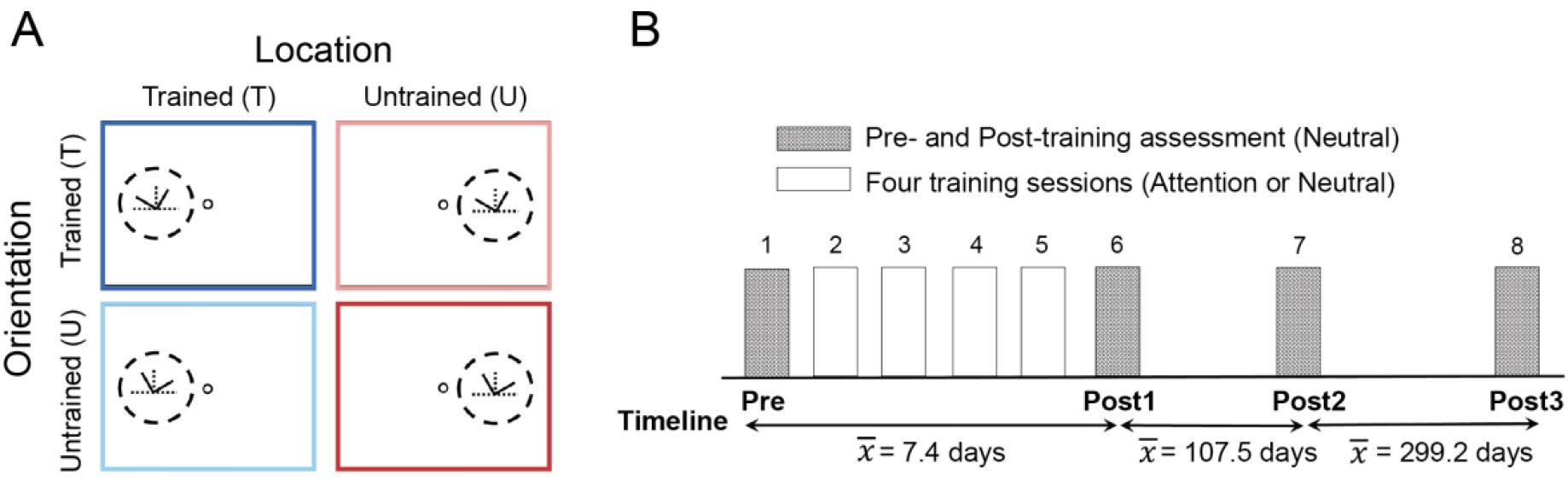
Illustration of the experimental conditions and protocol in the perceptual learning study. (**A**) Trained and untrained conditions in the testing sessions. The dark blue, light blue, light red, and dark red panels represent trained location/orientation (TL, TO), trained location/untrained orientation (TL, UO), untrained location/trained orientation (UL, TO), and untrained location/orientation (UL, UO), respectively. (**B**) Schematic illustration of the 6-day VPL experiment. Observers were tested on day 1 (Pre-test) and day 6 (Post-test 1), and were trained with a neutral cue or an attentional cue on days 2-5 depending on their assigned group. Observers performed an identical testing session 3-4 months (Post-test 2) and more than 1 year (Post-test 3) after completion of the VPL experiment to assess the duration of training effects of perceptual learning.

For both Attention and Neutral groups, we employed the method of constant stimuli during the testing (400 trials each) and the training (800 trials each) sessions. Observers’ performance was assessed for the five orientation offsets (**Supplementary Figs S1,S2** online), and 75% accuracy threshold was estimated by fitting a power function (for details, see Materials and Methods). Our training protocol was effective, as confirmed by significant learning in the Neutral group (**Fig 3A**, dark blue circles on sessions 1 vs. 6, t_8_=8.973, *p*<0.001, Cohen’s d=2.880, two-tailed, paired t-test). As expected, The Neutral group showed location and orientation specificity: Learning in the orientation discrimination task did not transfer to any of the three untrained conditions (**Fig 3A**, *p*=0.97, 0.19 and 0.15 for light blue, light red and dark red circles on sessions 1 vs. 6, respectively, two-tailed, paired t-tests).

**Fig 3.**
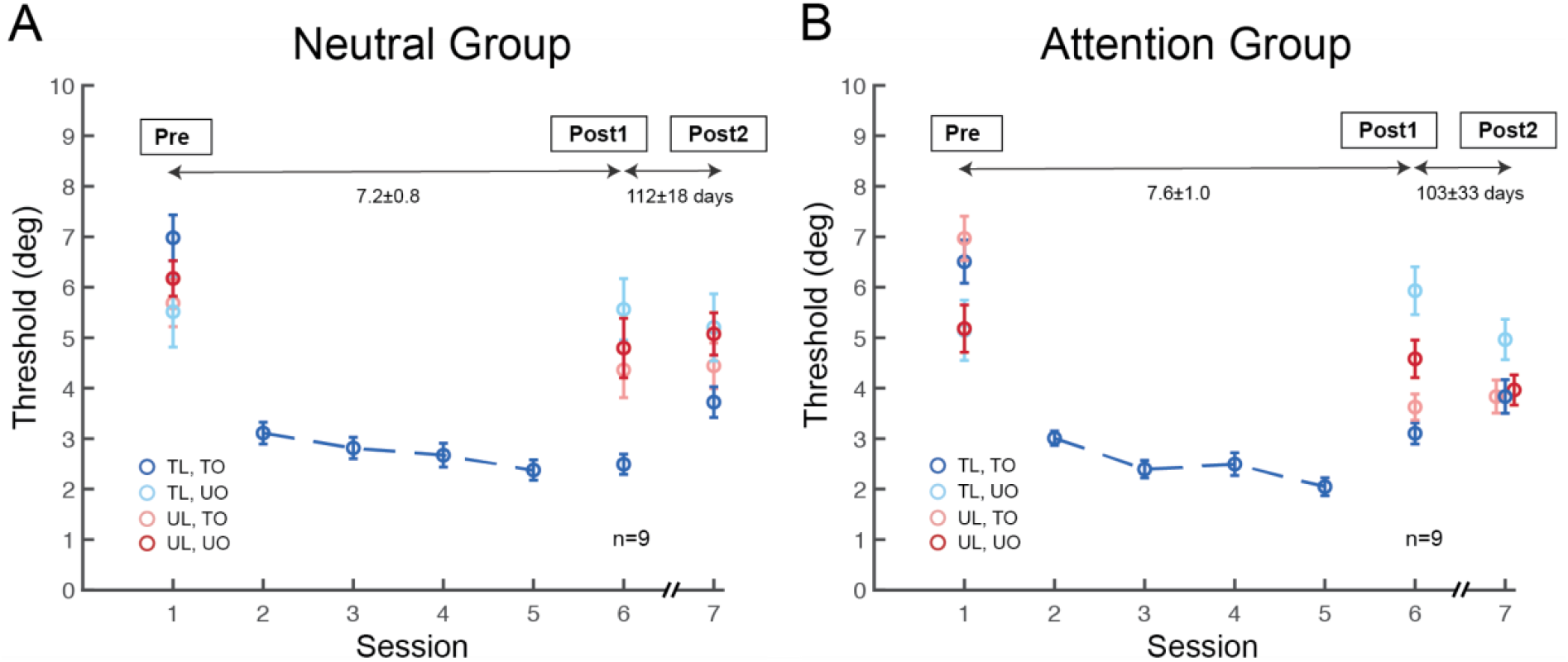
Spatial-transfer, but not feature-transfer, induced by feature-based attention in perceptual learning. (**A**) Session-by-session thresholds for orientation learning in the Neutral group. Performance improved for the trained feature at the trained location (dark blue circles), but not in the other three untrained conditions. The Neutral group exhibited both location- and feature-specificity in Post-test 1, as well as in Post-test 2. (**B**) Session-by-session thresholds for orientation learning in the Attention group. Similar to the Neutral group, observers retained improvement in the trained condition in both Post-test 1 and Post-test 2. (dark blue circles). Remarkably, training with FBA enabled complete learning transfer to the untrained location (light red circles), but not to the untrained orientation (light blue, dark red circles). Moreover, the improvement and location transfer induced by FBA persisted up to 3-4 months after training (dark blue, light red circles). Error bars represent ±1 within-subject SEM.

We next examined whether and how training with FBA affects VPL specificity. As in the Neutral group, observers in the Attention group showed significant learning in the trained condition (**Fig 3B**, dark blue circles on sessions 1 vs. 6, t_8_=6.336, *p*<0.001, Cohen’s d=2.647, two-tailed, paired t-test). The threshold differences between sessions 1 and 6 did not differ significantly between the two groups (t_16_=1.478, *p*=0.159, two-sample t-test), indicating comparable performance improvement in the trained condition. Likewise, performance in training sessions was similar between groups. A two-way ANOVA revealed a main effect of training (*F*(3,48)= 8.095, *p*<0.001), but no main effect of group or interaction between training and group (*F*<1). Importantly, unlike in the Neutral group, the Attention group showed complete learning transfer to the untrained location in the other hemifield (**Fig 3B**, light red circles on sessions 1 vs. 6, t_8_=5.225, *p*<0.001, Cohen’s d=1.706, two-tailed, paired t-test), with a comparable magnitude of performance change to the trained condition (**Fig 6B**, dark blue and light red bars, t_8_=0.617, *p*=0.555, two-tailed, paired t-test). However, learning did not transfer either to the untrained orientation (**Fig 3B**, light blue circles on sessions 1 vs. 6, t_8_= −0.784, *p*=0.456, two-tailed, paired t-test) or to the untrained orientation at the untrained location (**Fig 3B**, dark red circles on sessions 1 vs. 6, t_8_=0.757, *p*=0.471, two-tailed, paired t-test).

To further investigate our hypothesis that FBA training induces location transfer, we conducted a three-way mixed ANOVA with within-subject factors of location (trained vs. untrained) and training (Pre-test vs. Post-test 1), and a between-subjects factor of group (neutral vs. attention) using threshold values at the trained orientation (**Fig 4A**). There was a significant main effect of training (*F*(1,16)=87.621, *p*<0.001), indicating that performance became better at Post-test 1 than at Pre-test. Critically, there was a significant three-way interaction among location, training and group (*F*(1,16)=5.437, *p*=0.033). A two-way ANOVA (location x training) for each group revealed that for the Neutral group, there was a main effect of training (*F*(1,8)=38.355, *p*<0.001), and an interaction between location and training (*F*(1,8)=7.7284, *p*=0.024), indicating greater learning at the trained than the untrained location (*p*=0.027). For the Attention group, there was also a main effect of training (*F*(1,8)=49.512, *p*<0.001), but no interaction between location and training (*F*<1), indicating that the extent of learning was comparable at both the trained and untrained locations.

**Fig 4.**
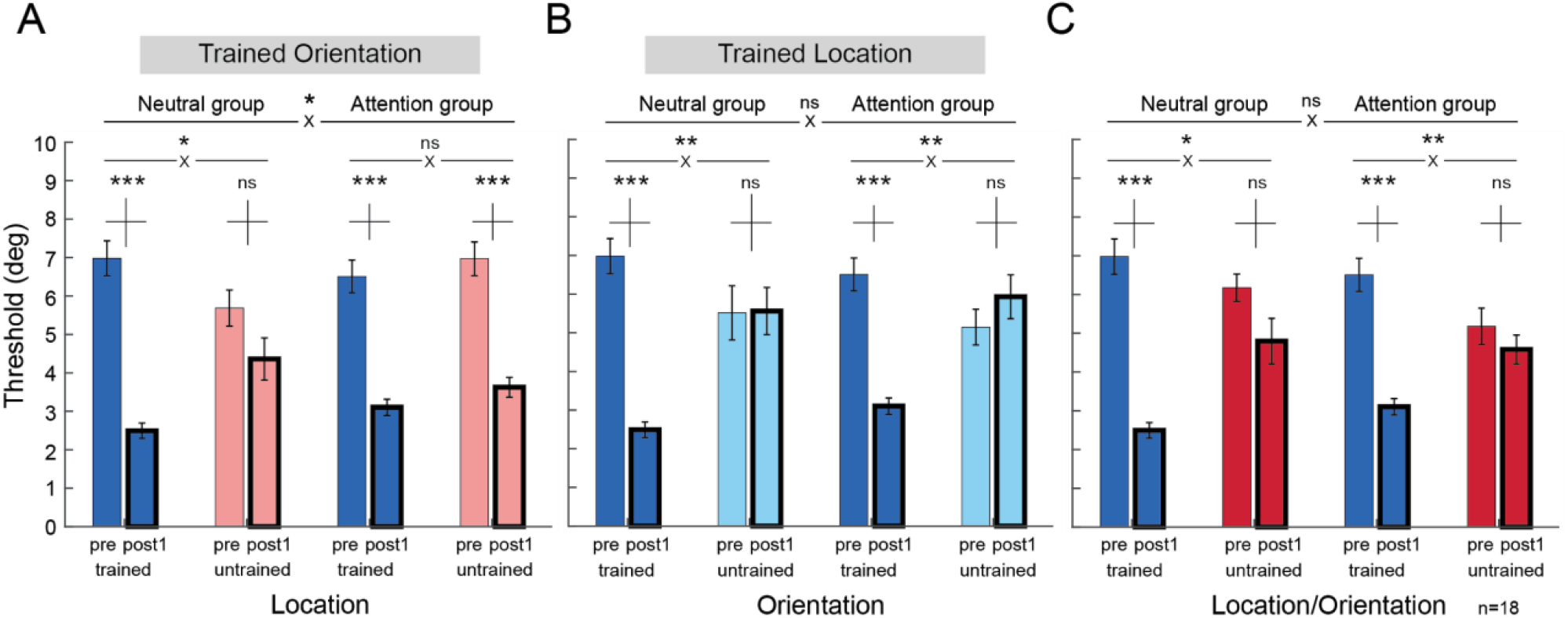
Threshold comparisons of Pre-test versus Post-test 1 between the Neutral and Attention groups. The trained condition was compared with the (**A**) untrained location, (**B**) untrained orientation, and (**C**) untrained location and orientation between the two groups. Learning transfer was found only in the untrained location in the Attention group (**A**), but not in the other untrained conditions (**B**,**C**) of either group (n=9 per group). * *p* < 0.05; ** *p* < 0.01; *** *p* < 0.001. Error bars represent ±1 within-subject SEM. Vertical bars above paired comparisons represent ±1 SEM for the mean threshold difference between Pre-test and Post-test 1.

There was an overall correlation between the pre-training threshold and the degree of improvement (*r*(80)=0.45, *p*<0.001). Thus we asked, could the pre-training threshold have affected the degree of transfer [60,61]? A two-way ANOVA revealed a main effect of condition (*F*(3,54)=3.022, *p*=0.037), but no main effect of group (*F*(1,18)=0.291, *p*=0.597) or interaction between condition and group (*F*(3,54)=1.853, *p*=0.148) on the Pre-test in the initial dataset (20 observers; Supplementary **Fig S3**). Despite no statistical significance, to prevent any possible confound due to a difference of the pre-training threshold for the trained orientation at the untrained location, we equated the pre-training thresholds in this condition by removing the observer with the lowest threshold in the Neutral group and the observer with the highest threshold in the Attention group; again, there was no significant difference between the two groups (**Fig 4A**, t_16_=1.244, *p*=0.232, two-sample t-test). Moreover, the pattern of results was the same when removing two observers from each group to further equate the threshold (Supplementary **Fig S4**): There was location transfer in the Attention group but not in the Neutral group.

We also conducted similar three-way mixed ANOVA to assess learning at the untrained orientation/trained location (**Fig 4B**), and at the untrained orientation/untrained location (**Fig 4C**). For each condition, there was a main effect of training (**Fig 4B**, *F*(1,16)=14.097, *p*=0.002; **Fig 4C**, *F*(1,16)=42.181, *p*<0.001), and a two-way interaction between condition and training (**Fig 4B**, *F*(1,16)=35.851, *p*<0.001; **Fig 4C**, *F*(1,16)=22.509, *p*<0.001), but no three-way interaction among condition, training and group (**Fig 4B**, *F*<1; **Fig 4C**, *F*<1). Therefore, learning did not differ between groups for these two untrained conditions. In sum, these results support our hypothesis that training with FBA unlocks location specificity, while preserving orientation specificity in VPL.

### Long-Term Retention of VPL Improvement and Transfer

Whereas it is well known that location transfer can be attained under certain experimental conditions [5,35,37,38,40] and that VPL improvement can last for months or even years [9,22–26], it is unknown how long learning transfer can last. To assess the duration of the observed VPL effects, we conducted a follow-up experiment. All 18 observers who completed the VPL training were recruited back 3-4 months after their Post-test 1 (*M*=107.5 days, *SD*=26.3 days), and completed the same testing session, which we refer to as Post-test 2.

In Post-test 2, learning of the orientation discrimination in the Neutral group (re-tested 112 ± 18 days after Post-test 1) remained robust in the trained condition (**Fig 3A**, dark blue circles on sessions 1 vs. 7, t_8_=5.947, *p*<0.001, Cohen’s d=1.772, two-tailed, paired t-test). Moreover, the Neutral group retained both location and orientation specificity (**Fig 3A**, *p*=0.81, 0.13 and 0.06 for light blue, light red, and dark red circles on sessions 1 vs. 7, respectively). Critically, for the Attention group (re-tested 103 ± 33 days after Post-test 1) not only the improvement remained at the trained condition (**Fig 3B**, dark blue circles on sessions 1 vs. 7, t_8_=4.593, *p*=0.002, Cohen’s d=1.841, two-tailed, paired t-test), but also at the transferred location (**Fig 3B**, light red circles on sessions 1 vs. 7, t_8_=4.301, *p*=0.003, Cohen’s d=1.654, two-tailed, paired t-test). We note that although performance for the untrained orientation at the untrained location in the Attention group seemed to improve from Post-test 1 to Post-test 2, this improvement did not reach significance and performance at Post-test 2 does not significantly differ from its Pre-test (dark red circles on sessions 1 vs. 7, t_8_=1.645, *p*=0.139).

To assess the training effects in a longer time scale, we conducted Post-test 3 one year after completion of training (**Fig 5**). Six observers in the Neutral group and 5 observers in the Attention group participated in Post-test 3, which took place 414 ± 34 days after their Post-test 1 (due to the COVID-19 pandemic, we were not able to recruit all observers back). Despite a long period of time, learning in the Neutral group (re-tested 408 ± 38 days after Post-test 1) remained significant in the trained condition (**Fig 5A**, dark blue circles on sessions 1 vs. 8, t_5_=3.497, *p*=0.009, Cohen’s d=1.174, one-tailed, paired t-test), but did not transfer to the three untrained conditions (**Fig 5A**, *p*=0.40, 0.28 and 0.26 for light blue, light red, and dark red circles on sessions 1 vs. 8, respectively; 1 of the 6 observers in the Neutral group lost improvement in Post-test 3). The Attention group (re-tested 420 ± 31 days after Post-test 1) retained learning at the trained condition (**Fig 5B**, dark blue circles on sessions 1 vs. 8, t_4_=3.02, *p*=0.02, Cohen’s d=1.993, one-tailed, paired t-test), as well as location transfer (**Fig 5B**, light red circles on sessions 1 vs. 8, t_4_=2.377, *p*=0.038, Cohen’s d=1.455, one-tailed, paired t-test). These results support our hypothesis that not only the improvement gained in both groups, but also the transfer induced in the Attention group is long lasting, up to over a year after training. We note that although performance for the untrained orientation at the trained location in the Attention group improved from Post-test 2 to Post-test 3, performance at Post-test 3 still does not significantly differ from its Pre-test (light blue circles on sessions 1 vs. 8, t_4_=0.518, *p*=0.632).

**Fig 5.**
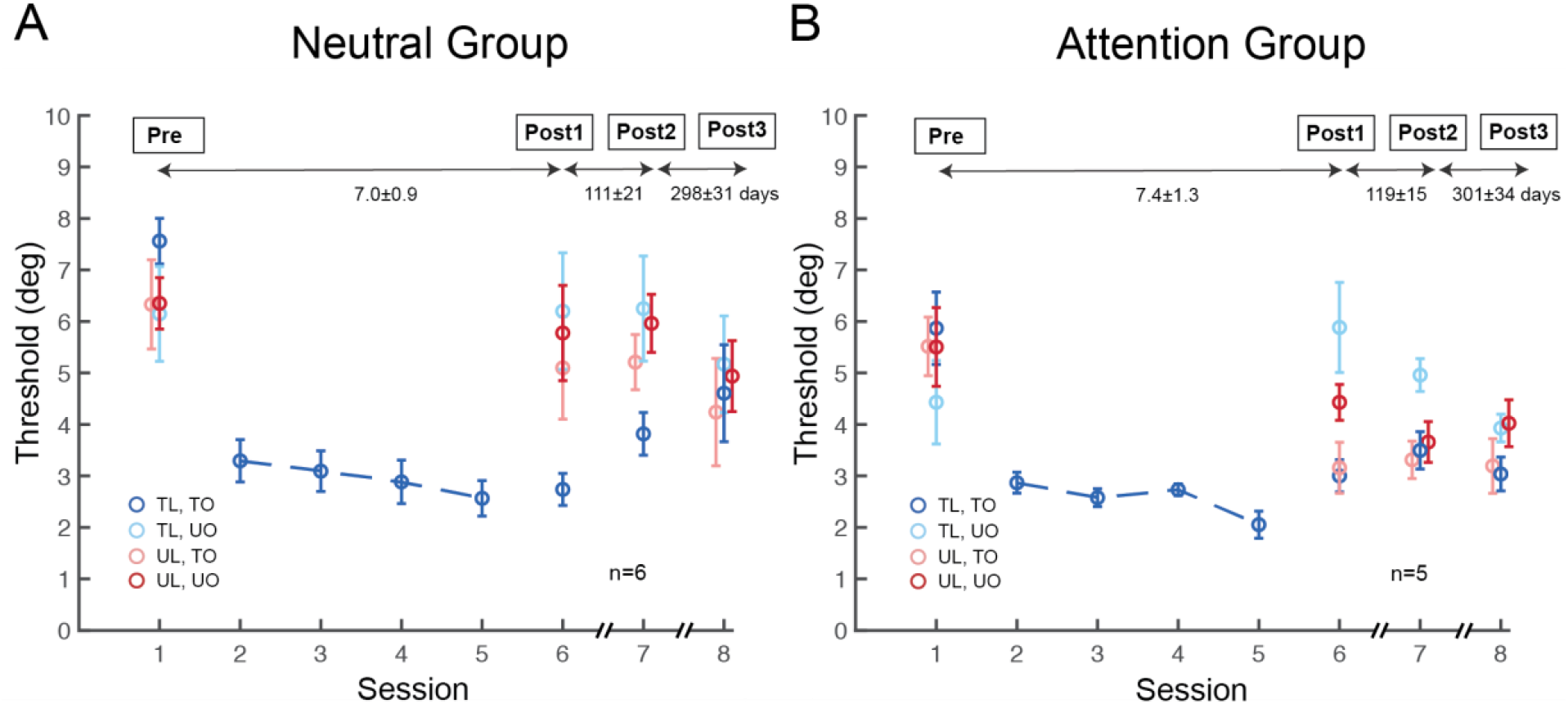
Session-by-session thresholds of Pre-test, Post-test 1, Post-test 2, and Post-test 3 in the Neutral and Attention groups. 6 observers from the Neutral group and 5 observers from the Attention group were re-tested 1 year after the completion of training. Similar to results in Post-tests 1 and 2, the perceptual improvement in both groups and location transfer in the Attention group were preserved. Error bars represent ±1 within-subject SEM.

We calculated observers’ Mean Percent Improvement (MPI) to analyze the magnitude of performance changes between the Neutral and Attention groups for individual conditions, and none of the comparisons was significant (*p*>0.05). To further compare performance changes at the post-tests between the groups, observers’ MPI was assessed between Pre-test versus Post-test 1, Pre-test versus Post-test 2 and Pre-test versus Post-test 3 (**Fig 6**). In Post-test 1, the Neutral group exhibited significant improvement, MPI= 63.2 ± 10.7% only at the trained condition (**Fig 6A**, dark blue bar, t_8_=20.035, *p*<0.001, Cohen’s d=9.444; light red bar=20.3 ± 9.1%, t_8_=1.279, *p*=0.237; light blue bar= −14.1 ± 13.7%, t_8_= −0.684, *p*=0.514; and dark red bar=21.5 ± 8.6%, t_8_=1.716, *p*=0.125, two-tailed, paired t-tests). The Attention group exhibited significant, comparable amount of improvement both in the trained condition and at the untrained location for the trained orientation (**Fig 6B**, dark blue bar=51.1 ± 10.9%, t_8_=8.273, *p*<0.001, Cohen’s d=3.9; light red bar=45.3 ± 12.3%, t_8_=7.112, *p*<0.001, Cohen’s d=3.353, two-tailed, paired t-tests). Such transfer did not occur either at the untrained orientation for the trained location (**Fig 6B**, light blue bar= −37.2 ± 22.5%, t_8_= −0.564, *p*=0.266), or at the untrained orientation and untrained location (**Fig 6B**, dark red bar= −4.9 ± 15.1%, t_8_= −0.105, *p*=0.829).

**Fig 6.**
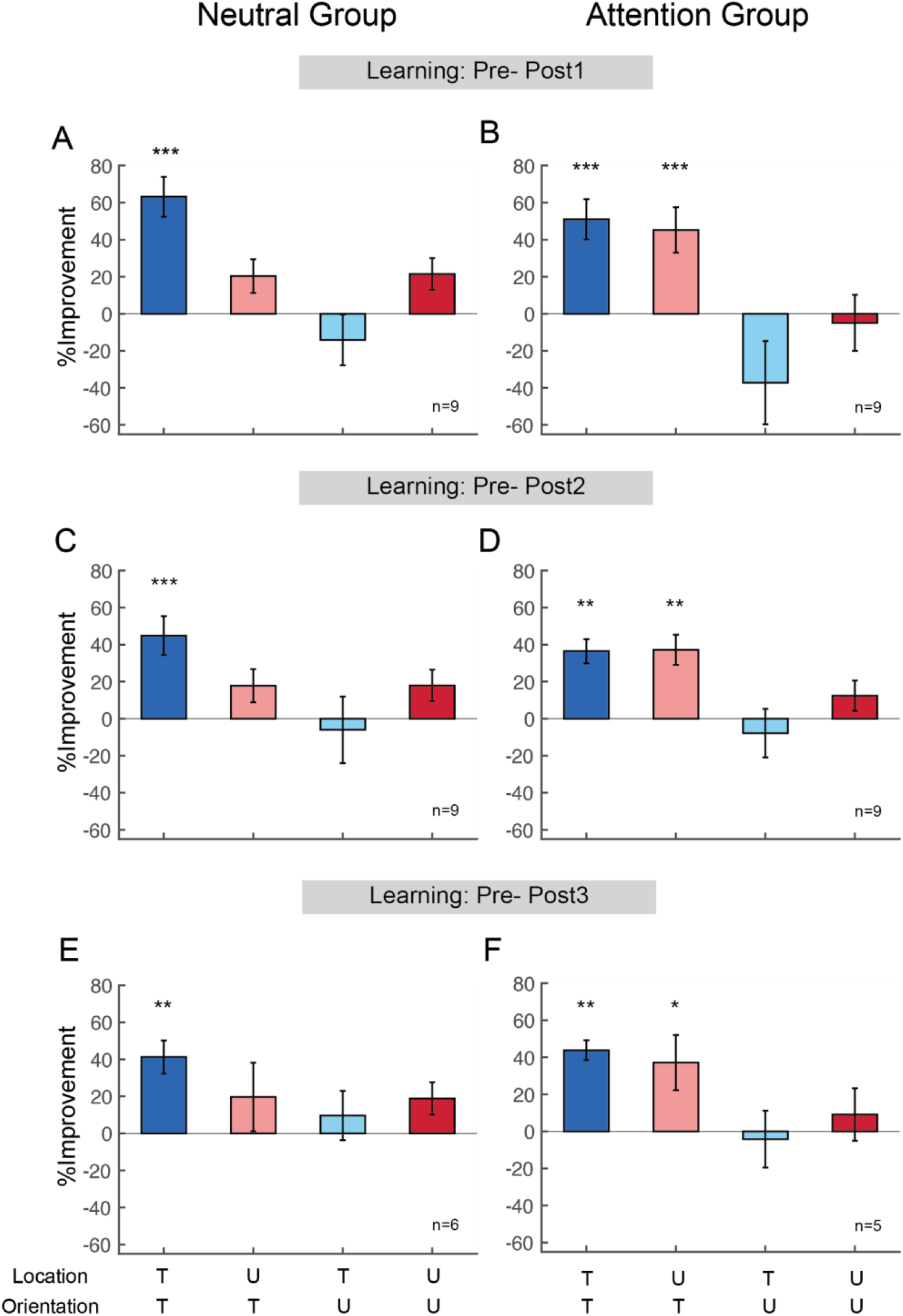
Performance changes at Post-test 1, Post-test 2 and Post-test 3 between the Neutral and Attention groups. (**A, B**) Performance changes between Pre-test and Post-test 1. The Neutral and Attention group showed significant learning in the trained condition (dark blue bars in **A**, **B**). Contrary to the Neutral group, training with FBA unlocked location specificity (light red bar in **B**), while preserving orientation specificity (light blue, dark red bars in **B**). (**C, D**) Performance changes between Pre-test and Post-test 2. Consistent with the results in Post-test 1, the improvement in both groups (dark blue bars in **C**, **D**) and location transfer in the Attention group (light red bar in **D**) were preserved 3-4 months after completion of training. (**E, F**) Performance changes between Pre-test and Post-test 3. The improvement in both groups (dark blue bars in **E**, **F**) and location transfer induced by FBA (light red bar in **F**) persisted longer than 1 year after training. * *p* < 0.05; ** *p* < 0.01; *** *p* < 0.001. Error bars represent ±1 within-subject SEM.

In Post-test 2, learning at the trained condition remained for both groups (**Figs 6C**, dark blue bar=44.8 ± 10.5%, t_8_=6.425, *p*<0.001, Cohen’s d=3.029; **6D**, dark blue bar=36.5 ± 6.5%, t_8_=4.312, *p*=0.001, Cohen’s d=2.033, one-tailed, paired t-tests). Importantly, in the Attention group, the improvement was retained at the untrained location for the trained orientation (**Fig 6D**, light red bar=37.2 ± 8.1%, t_8_=4.47, *p*=0.001, Cohen’s d=2.107). Although the amount of improvement decreased slightly from Post-test 1 to Post-test 2 (**Figs 3A,B**, dark blue circles on sessions 6 and 7; **Fig 6**, dark blue bars in **A,B** vs. **C,D**), the pattern remained: training with FBA induced complete location transfer both in Post-test 1 and Post-test 2 (**Figs 6B,D**).

In Post-test 3, both groups still retained improvement at the trained condition (**Figs 6E**, dark blue bar=41.2 ± 9.0%, t_5_=3.859, *p*=0.006, Cohen’s d=2.223; **6F**, dark blue bar=43.8 ± 5.4%, t_4_=4.27, *p*=0.006, Cohen’s d=2.707, one-tailed, paired t-test). Furthermore, in the Attention group learning transfer to the untrained location remained (**Fig 6F,** light red bar=37.1 ± 14.9%, t_4_=2.302, *p*=0.041, Cohen’s d=1.456, one-tailed, paired t-test). There was no transfer in the other untrained conditions in either the Neutral group (**Fig 6E**, t_5_=0.732, *p*=0.497 for light red bar, t_5_=0.676, *p*=0.529 for light blue bar and t_5_=1.132, *p*=0.309 for dark red bar) or the Attention group (**Fig 6F**, t_4_= −0.178, *p*=0.868 for light blue bar and t_4_=0.362, *p*=0.736 for dark red bar).

Taken together, our results show a perceptual benefit of FBA on an orientation discrimination task and reveal a remarkable spatial-transfer of learning induced by FBA. Moreover, the perceptual improvements in the trained condition in both groups and the learning transfer to an untrained location in the Attention group were preserved for over a year, indicating robust, long-term benefits of training with FBA in VPL.

## Discussion

This study reveals how training with FBA benefits human VPL. We first confirmed that FBA improved accuracy in an orientation discrimination task. Next, manipulating FBA during training induced spatial-transfer, but not feature-transfer, in VPL. Our findings that FBA unlocks location specificity, while preserving feature specificity, are consonant with the psychophysical and neural evidence showing that FBA effects are independent of the location of the attended stimuli [47–58], and expand our understanding of FBA’s global modulation from human visual perception to VPL. Critically, the perceptual improvements and generalization gained from our training protocol persisted for over a year, revealing that FBA induces long-term benefits.

It has been established that VPL transfer is related to particular training protocols [33,35–38,46,60,61]. Its presence and degree is influenced by numerous experimental factors, including length of training [33], stimulus precision [35,60], variability of training stimuli [61], initial threshold and the amount of learning [62,63]. Retinotopic specificity is preserved with high stimulus precision during training [35] or in a transfer task [60]. We note that in the present study we controlled for the amount of training, stimuli precision and variability (using the method of constant stimuli, with 5 levels of orientation offset) and equated the initial thresholds of the trained condition between the Neutral and Attention groups. Although the method of constant stimuli, due to stimulus variability, could lead to transfer, we found location and orientation specificity in the Neutral group. Furthermore, it is likely that the internal representation of the reference orientations becomes less noisy throughout training [64], but such effect would have been the same at the trained and untrained locations/orientations. Therefore, our training procedure in the two groups could have not been differentially subjected to any of the aforementioned factors that can impact VPL specificity. Moreover, our experimental design ensured that FBA deployment during training was the critical influence determining robust location transfer. The cues only differed during the training days for the Neutral and Attention groups, but both groups performed the pre- and post-test sessions with a neutral cue. Analogous designs have been used to manipulate and isolate the role of covert spatial attention in learning acquisition [39] and in location transfer [5,27,40].

In addition to the remarkable location-transfer induced by FBA, we also observed a slight (not statistically significant) suppression in performance at the untrained features in the Attention group (**Figs 6B,D**, light blue bars). This observation is consistent with studies showing FBA enhancement of the target and suppression of nearby features [65–68]. The ‘feature-similarity gain model’ highlights the similarity between the attended features and the neurons’ preferred features. Enhancement takes place when the attended stimulus matches the neurons’ preference, and suppression when the attended feature is dissimilar to the neurons’ preferred feature [52,69].

Regarding the neural basis underlying the behavioral improvement in orientation discrimination tasks, it has been shown that training alters the tuning profiles of populations of orientation-selective neurons in the visual cortex [7,70,71]. A monkey single-unit recording study reported that orientation learning led to an increase in neuronal selectivity in V1, with steeper tuning functions for the neurons most sensitive to the trained orientation [7]. Likewise, a human fMRI study revealed that extensive training can refine neural representations in the occipital cortex (V1-V4), even in the absence of gross changes of BOLD responses after training [70]. The training-induced changes manifested as specifically enhanced neural representation of the trained orientation at the trained location, in agreement with specificity of perceptual learning in their behavioral task.

What mechanism could drive location transfer induced by FBA in VPL? Reweighting models have been proposed to account for specificity and transfer in VPL [72–75]. To explain transfer across retinotopic locations, the Integrated Reweighting Theory builds a multi-level learning system that incorporates higher-level, location-independent representations with lower-level, location-specific representations, which are both dynamically modified in VPL [74]. The performance improvement results from pruning weights on untrained orientations, and amplifying weights on relevant, trained orientations to the decision unit [76,77]. Whereas location transfer is mediated by reweighting the broadly tuned location-independent representations, specificity arises from reweighting the narrowly tuned location-specific representations [74]. At the performance level, FBA improves visual perception in a way consistent with boosting the gain and sharpening the tuning of neuronal population responses to the attended features [78,79], consistent with neurophysiological [69] and neuroimaging [70,80] findings. At the learning level, deploying FBA during perceptual training is likely to shape the neural circuits through increasing the weights between location-independent representations and the decision unit, thus mediating location transfer in VPL.

Spatial attention also facilitates location transfer in VPL [5,27,40]. However, the mechanisms underlying this transfer, which seems counter-intuitive to the localized effect of spatial attention, remain unclear. It has been speculated that the short-term improvement of sensory signals due to spatial attention may enable a higher level learning mechanism [5,27,40]. The Reverse Hierarchy Theory [37,86] predicts specificity in difficult tasks in which training modifies low-level, location- and feature-specific units, and transfer in easy tasks in which training-induced modifications are at high-level, broadly-tuned units. Accordingly, because covert spatial attention enhances sensory processing and improves performance [45,81], making tasks less difficult, learning may rely more on high-level units and facilitate transfer. Given that the stimulus parameters, task and training days of the current study differ from those of spatial attention, it is not possible to directly compare their transfer effects. Future research with a constant experimental design is required to compare the degree of location transfer induced by spatial attention and FBA.

Elucidating the mechanisms underlying specificity and transfer has become a central focus of the VPL field. There is substantial evidence supporting that specificity reflects plasticity in low-level brain areas [7,29–32], but learning-related neuronal changes are not only confined to the primary sensory areas [71,82–84]. Research indicating that specific learning effects can arise from top-down influences [85–87], or can be accomplished by changes of readout weights in decision areas, highlight the importance of higher-level brain areas involved in VPL [73,74,76,77]. These studies, as well as the current findings, suggest that VPL involves low-level representations, higher level representations, read-out, attention, and decisions [88], and that changes in one or multiple brain systems could determine the degree of behavioral learning transfer.

Findings from VPL studies have translational implications for improving visual expertise and clinical rehabilitation. To promote VPL generalization, researchers must develop efficient protocols that overcome specificity to maximize training benefits. The present study provides an important step in optimizing visual training protocols to promote learning generalization, which could have translational value for developing training tools. The effectiveness of FBA training with a special population provides converging evidence for the usefulness and potential of this approach. A recent study has shown that cortically blind patients who trained with FBA could restore performance in a fine-direction discrimination task, whereas those patients who trained without FBA could not [89].

To conclude, we have implemented an elegant, well-controlled design to assess whether and to which extent FBA affects the degree of location and feature specificity, which are hallmarks of VPL. The pronounced, long-lasting training benefits we observed reveal FBA as an effective tool to generalize learning across locations over a long time scale. Furthermore, these findings can inform models and theories that link visual learning to plasticity across multiple cortical areas and can shed light on our understanding of the underlying neural mechanisms of attention-induced transfer in human learning.

## Materials and methods

### Observers

Eighteen (12 females; mean=26.6 ± 6.4 years old) and other 20 (13 females; mean=23.6 ± 4.6 years old) human observers who had normal or corrected-to-normal vision participated in Experiments 1 and 2, respectively. In Experiment 1, we did not include people whose data quality fell below our criteria (overall 75% accuracy). In Experiment 2, one observer was excluded from each group for analysis to equate pre-training thresholds for the untrained location, trained orientation condition across groups. We note that all reported results are the same when we take all 10 observers per group into account (Supplementary **Fig S3**), or remove two from each group for a further threshold equation (Supplementary **Fig S4**). The 18 remaining naïve observers were equally distributed into two groups – Neutral (5 females; mean= 21.2 ± 1.8 years old) or Attention (7 females; mean= 24.0 ± 4.6 years old). The experimental protocols were approved by the University Committee on Activities Involving Human Subjects of New York University, and all research was performed in accordance with relevant guidelines/regulations. Informed consent was obtained from all observers.

### Apparatus

The stimuli were presented using Psychophysics Toolbox [90,91] for MATLAB (The MathWorks, Natick, MA) on an iMac computer with a 21” gamma-corrected Sony GDM-5402 CRT monitor with resolution of 1280 x 960 pixels and a refresh rate of 100Hz. An infrared eye tracker system Eyelink 1000 (SR research, Kanata, Ontario, Canada) and a chin rest and head rest were used to ensure eye fixation at the center of the display throughout each trial in the experimental sessions. The viewing distance was 57 cm, and all experiments were performed with a gaze-contingent display in which the eye-tracker enabled new trials to start only once observers had fixated at the center (within a 2° radius fixation window). If an eye-movement outside of this window was detected at any point after the trial started, then that trial was aborted and added to the end of each block (~5% of the trials).

### Stimuli

In each trial, the stimulus was a single Gabor patch (Gaussian windowed sinusoidal gratings) subtending 2° of visual angle and presented at 5° eccentricity on a grey background. The Gabor had spatial frequency of 4 cpd, standard deviation of 2λ, and contrast of 0.64. To assess five different difficulty levels, there were five offsets (2°, 4°, 6°, 8°, and 10°) that were either clockwise or counter-clockwise from reference angles. We used four reference orientations, which indexed different features in this task, and at the beginning of each block two of the references (either reference combination 1 of 30°/120°, or combination 2 of 60°/150°) were presented simultaneously (**Fig 1A**). The neutral cue consisted of a pair of leftward and rightward arrowheads flanking the fixation dot, each starting 0.6° from the fixation point and composed of two 0.5°-long x 0.12°-wide black lines 92° apart. The attentional cue was either a leftward arrowhead indicating a reference angle of 30° or 60°, or a rightward arrowhead indicating 120° or 150°, depending on the reference combination for that block (**Fig 1B**). For all blocks in Experiment 1, and the testing sessions of Experiment 2, the feedback was a 1°-long x 0.06°-wide line on top of a white fixation dot (radius 0.15°) presented at the reference angle of the just-perceived stimulus, to remind observers of the exact reference orientations. For all training sessions of Experiment 2, the feedback was given at the fixation dot indicating trial accuracy.

### Orientation discrimination task

Each trial began with a 400-ms fixation period followed by a 200-ms neutral or attentional cue (**Fig 1A**). After a 400-ms ISI, the stimulus was presented for a single 200-ms interval, and the observer’s task was to judge whether the orientation of the stimulus was clockwise or counter-clockwise to the closest reference orientation by pressing labeled keys “/” or “\” on the keyboard, respectively. The temporal parameters ensured that observers had time to deploy FBA [48]. Two reference lines were shown to observers before each block, but never appeared on the screen during stimulus presentation, so observers were encouraged to use their internal representation of the reference orientations to perform the discrimination. Observers had 4 s to indicate their answer by a key-press, and then received a 300-ms feedback line flashing green for correct responses, or red for incorrect responses. There was a 1-s inter-trial interval.

### Practice

In Experiments 1 and 2, observers performed 4 practice blocks (20 trials each) of the orientation task, with reference combinations (30°/120° or 60°/150°) and locations (left or right) counterbalanced and a 10° offset between targets and references. The criterion was 70% accuracy before proceeding to the main task. Additionally, on day 1 of Experiment 2, observers completed 40 trials of a simple color-discrimination task before the orientation practice, to familiarize themselves with the procedure and timing, and to reduce procedural learning during the perceptual learning experiment.

### Experiment 1

This experiment consisted of a single session of the orientation discrimination task. Observers performed 800 trials, equally distributed in four 200-trial sections that corresponded to either a neutral (N) or an attentional (A) cue. The four sections were administered in N-A-N-A or A-N-A-N counterbalanced order. The 200 trials in each section were divided into 4 blocks, each corresponding to a different condition (i.e., stimuli on the left or right, and reference orientations of 30°/120° or 60°/150°), and 50 trials (5 repetitions of each the 5 offset-sizes, and each of the two reference angles) were randomized per block. To use the attentional cue, participants were instructed to deploy their attention to a particular feature (a reference orientation) indicated by the cue before the stimulus presentation (**Fig 1B**). Given the nature of simultaneous features in our design, and that no explicit reference was shown during orientation discrimination, this task was difficult even for experienced observers.

### Experiment 2

This was a six-day perceptual learning experiment. Observers were tested at 5° eccentricity on left or right horizontal meridian for each of the reference combinations −30°/120° or 60°/150°– on their first and sixth days and completed 4 training sessions on days 2-5 (**Figs 2A,B**). All 6 sessions were performed at the same or a similar time across the average time frame of 7.4 days (SD = 0.9 day), with ≤ 2 days between consecutive sessions.

The testing sessions consisted of 400 trials, all presented with a neutral cue, equally distributed between four different conditions (i.e., stimuli on the left or right, and reference orientations of 30°/120° or 60°/150° counterbalanced across observers; **Fig 2A**). Each of the 4 conditions contained two blocks of 50 trials (5 trials per offset size and reference angle). The order of the eight blocks was randomized. During the training sessions, observers performed one condition for 800 trials, with 400 trials for each reference orientation, with a neutral or attentional cue depending on the group assignment. The 800 trials were split into 16 blocks of 50 trials (5 trials per offset size and reference angle), with short breaks between blocks and a 5-min break in the middle of the session. In addition, to assess how long the training effects would last, all observers from the two groups were recruited back 3-4 months after completion of the six-day experiment and asked to perform the same testing session (i.e., Post-test 2; **Fig 2B**). Moreover, 6 observers from the Neutral group and 5 observers from the Attention group were re-tested 1 year after completion of the six-day experiment (i.e., Post-test 3; **Fig 2B**). Due to the COVID-19 pandemic, we could not recruit the rest of the observers.

## Data analysis

Performance in the orientation discrimination task was measured using the method of constant stimuli (across five orientation offsets). Mean observer performance was plotted as psychometric curves across five orientation offsets, then fitted by a power function with R^2^ to indicate the quality of fit (**Figs 1C,S1,S2**).

Threshold in the VPL study was estimated by a power function (*f*(*x*) = *ax^n^*, where a is a constant and n is a real number) where observers achieved 75% accuracy.

In Experiments 1 and 2, repeated measures ANOVAs were performed in MATLAB to assess statistical significance. A two-way ANOVA was conducted to assess the effects of attention and orientation offsets on accuracy in Experiment 1. For Experiment 2, we conducted a three-way ANOVA with within-subject factors of condition (trained vs. untrained) and training (Pre-test vs. Post-test 1), and a between-subjects factor of group (neutral vs. attention) using threshold values to examine potential interactions between the two groups. When a three-way interaction was found, a two-way ANOVA (condition x training) was conducted to assess the threshold changes after learning for each group (**Fig 4**). Paired t-tests were used to assess the threshold changes before and after training for each condition (**Figs 3,4,5**), and the performance changes for conditions within each group (**Fig 6**). The performance changes at post-tests were calculated as (Threshold_pre_ - Threshhold_post_) / Threshold_pre_ for each observer and represented as Mean Percent Improvement (MPI) in **Fig 6**. Error bars in all figures represent ±1 within-subject SEM [92], which takes into account individual variability by subtracting the group mean from each individual’s value.

## Acknowledgements

This research was supported by National Eye Institute EY016200 and EY027401 to MC. We thank members of the Carrasco Lab for their helpful comments.

## Author contributions

S.-C.H. and M.C. conceived and designed the experiments. S.-C.H. performed the experiments and analyzed data. S.-C.H. and M.C. wrote the paper.

## Competing interests

The authors declare that no competing interests exist.

## Notes

### Competing Interest Statement

The authors have declared no competing interest.

