## supplemental for "Feature-based attention enables robust, long-lasting location transfer in human perceptual learning"

**SUPPLEMENTAL FIGURES**

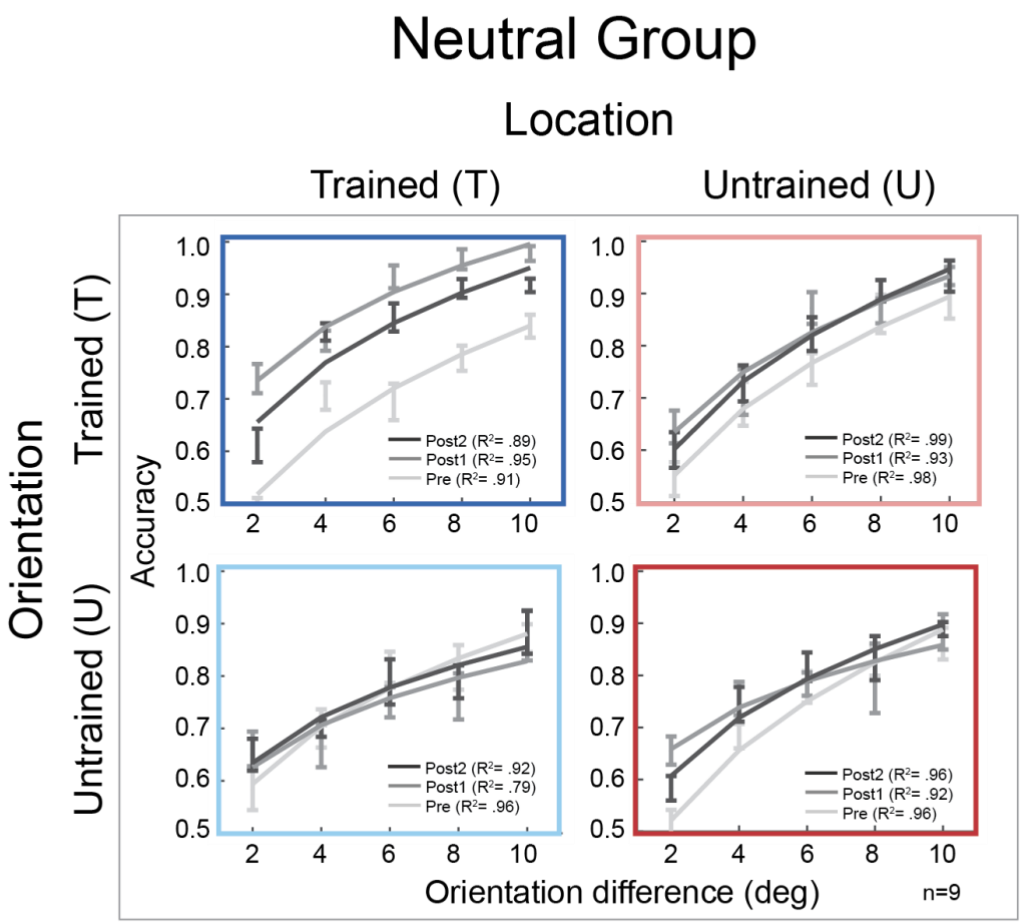


**Fig S1. Psychometric curves from the VPL study in the Neutral group. Related to Fig 3.** Results of Pre-test (light gray), Post-test 1 (dark gray), and Post-test 2 (black) for the trained and untrained conditions. Observers in the Neutral group exhibited both location and feature specificity in Post-test 1, as well as in Post-test 2. Error bars represent ±1 within-subject SEM.

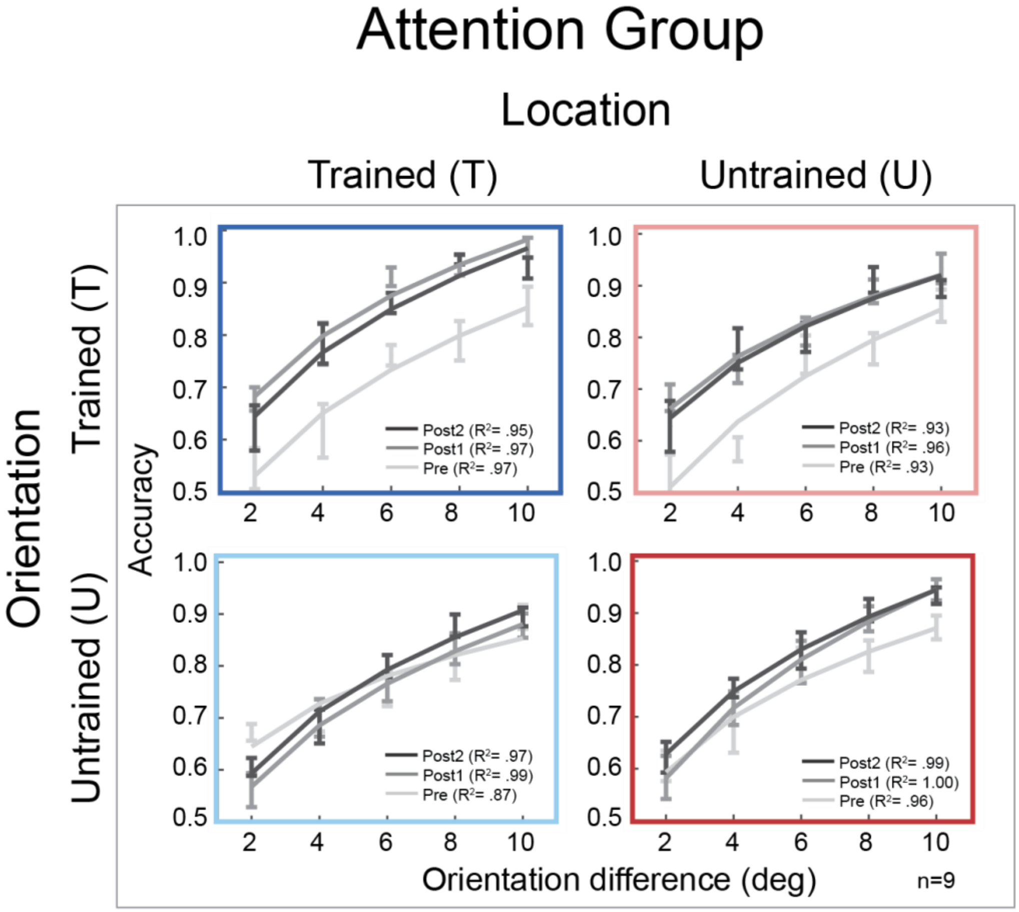


**Fig S2. Psychometric curves from the VPL study in the Attention group. Related to Fig 3.** Results of Pre-test (light gray), Post-test 1 (dark gray), and Post-test 2 (black) for the trained and untrained conditions. Similar to the Neutral group, observers showed improvement in the trained condition in both Post-test 1 and Post-test 2 (dark blue panel). Remarkably, training with FBA enabled complete learning transfer to the untrained location (light red panel), but not to the untrained orientation (light blue, dark red panels). Moreover, the improvement and location transfer persisted up to 3-4 months after training (dark blue, light red panels). Error bars represent ±1 within-subject SEM.

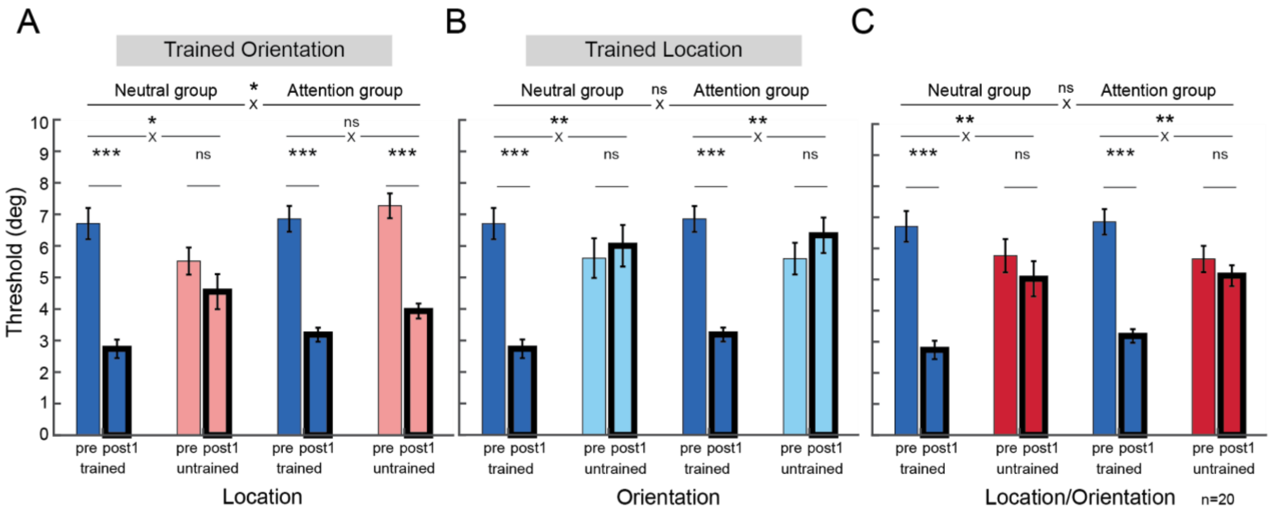


**Fig S3. Threshold comparisons of Pre-test versus Post-test 1 between the Neutral and Attention groups. Related to Fig 4.** The trained condition was compared with the (**A**) untrained location, (**B**) untrained orientation, and (**C**) untrained location and orientation between the two groups. Learning transfer was found only in the untrained location in the Attention group (**A**), but not in the other untrained conditions (**B**,**C**) of either group (n=10 per group). * $p < 0.05$; ** $p < 0.01$; *** $p < 0.001$. Error bars represent ±1 within-subject SEM.

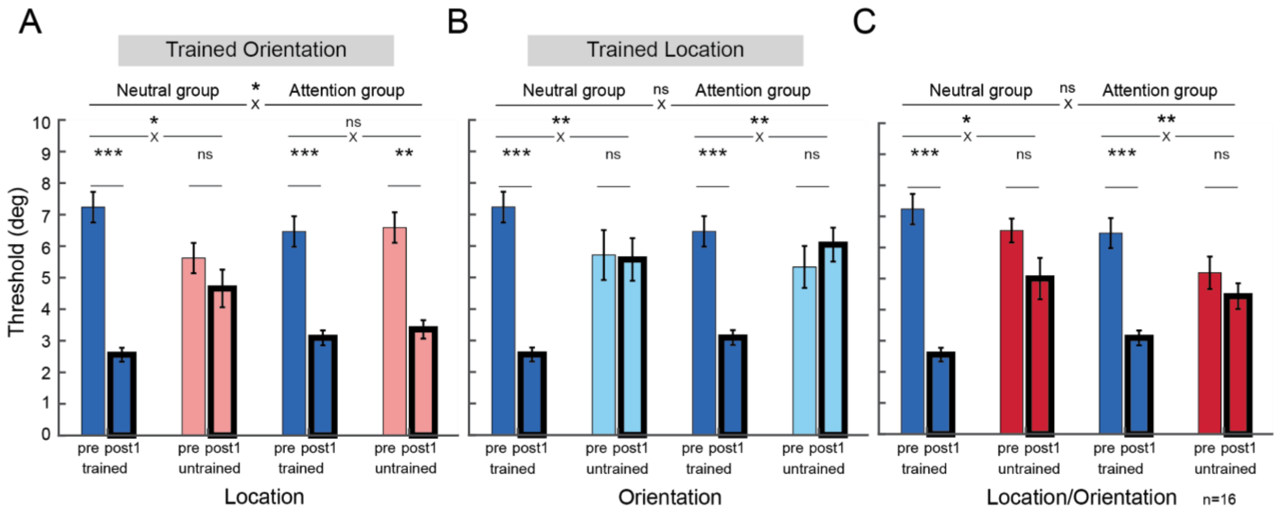


**Fig S4. Threshold comparisons of Pre-test versus Post-test 1 between the Neutral and Attention groups. Related to Fig 4.** The trained condition was compared with the (**A**) untrained location, (**B**) untrained orientation, and (**C**) untrained location and orientation between the two groups. Learning transfer was found only in the untrained location in the Attention group (**A**), but not in the other untrained conditions (**B**,**C**) of either group (n=8 per group). * $p < 0.05$; ** $p < 0.01$; *** $p < 0.001$. Error bars represent ±1 within-subject SEM.
